# Cooling Induced by Uphill Energy Transport in Plant Photosystems

**DOI:** 10.1101/146548

**Authors:** Koel Sen, Abhishek Bhattacharya, Santiswarup Singha, Maitrayee Dasgupta, Anjan Kr Dasgupta

## Abstract

The uphill energy transfer in photosystems implies input energy at higher wavelength leading to energy output at lower wavelength. Briefly, energy is uphill transported from photosystem I (PSI) to photosystem II (PSII), the latter having a lower wavelength emission. This uphill energy transport involves absorption of thermal energy from the surroundings. While such cooling effects have been reported in laser systems we report for the first time a white light driven cooling in thylakoid suspension. The cooling of the surrounding medium by appropriate illumination was illustrated using thermal measurements. Again cooling is inhibited by agents like 3-(3,4-Dichlorophenyl)-1,1-dimethylurea,that block the linear electron flow between the photocenters, implying a dependence of the cooling on interplay between such centers. Furthermore, it is possible to modulate the cooling pattern by addition of external agents like nanopaticles, some favoring further cooling (e.g., Ag nanoparticle) and some like Au or chlorophyll nanoparticles, showing insignificant or even reverse trends. Interestingly, the cooling is invariably associated with the 77K spectra of the thylakoid suspension. With reference to the dark control, an agent causing cooling always increases PSII to PSI ratio and vice versa i.e.,the uphill energy transport. Importantly, the cooling effect, apart from its import role in plant physiology can be exploited artificially for energy saving in post-harvest or food preservation.

## Introduction

Photosynthetic apparatus is the only natural photo electric engine sophisticated enough to convert light energy with an energy transduction efficiency of near unity (Scholes et al., 2011). One consensus that has been put forward to explain this high efficiency is non-classicality of the process(OReilly and Olaya-Castro, 2014). The features necessary to identify the non-trivial quantum phenomena explaining the coherent behaviour even at room temperature has recently been debated (Ishizaki and Fleming, 2012, Kassal et al., 2013, Lee et al., 2007, Norris, 1976, Panitchayangkoon et al., 2010, Ritz et al., 2010, Scholes, 2010).

An additional question namely, whether higher efficiency of photon capture implies absence of any thermal exchange with the surroundings, remains unanswered. The structural hierarchy of available photosynthetic materials consists of chlorophyll molecules, derivatives of chlorophyll, and other accessory pigments pigment-containing membrane spanning proteins, such as photosystem I and II represented conventionally as PSI, PSII(Ben-Shem et al., 2003, Duysens, 1964, Hipkins and Baker, 1986, Jennings et al., 2003, Lea et al., 1999).

Normally, the photosynthetic reaction centres embedded as protein complex and are present as parts of a double membrane envelope constituting of the thylakoid membrane. When exposed to illumination favouring either PSII or PSI, the system distributes the excitation such that the light-limited photo-system receives more energy at the expense of the other (Bonaventura and Myers, 1969). This light induced change in excitation transfer efficiency is slow in room temperature and fast in liquid nitrogen temperature, but the underlying mechanism is thought to be the same (Murata, 1969). Sen et al., have recently demonstrated that energy distribution between photosystems PSI and PSII oscillates with the progress of prolonged and continuous white light irradiation at varying intensities (Sen et al., 2014) implying a coupling between the energy transfer mediated by the respective photon capture complexes. This inference when compared with observations made in a recent paper that demonstrated that energy can migrate from high wavelength absorbing red chlorophylls to the reaction center via a slow thermally activated transfer based on the excitation equilibrium model (Jennings, Zucchelli, 2003)suggests that thermal energy coupling between the PSI, PSII energy transfer is also likely. While the cited works confirm coupling between the photocentres, the exact mechanism of the uphill energy transfer from PSI to PSII, the former emitting at higher wavelength (lower energy) remains unclear. Structurally the phenomenon is explained by existence of the gap chlorophylls which bridge the far red chlorophylls with the bulk chlorophylls of the inner core complex and biophysically by a slow equilibration phase in the excitation dynamic within the PSI-LHCI network. (Palacios et al., 2002). But the exact mechanism by which the energy I actually transferred, and such transfer is related to the thermodynamics of the photon capture process remains a grey area, fertile for further research. Interestingly, the basic photon capturing molecules present in the photo-systems are fluorescent in nature (Rosenqvist and van Kooten, 2003), and such property involves three attributes, namely excitation, emission and a non-radiative energy transfer (Frster, 1959). The thermodynamic puzzle a natural photo-system poses, is not only in the efficient conversion of photon energy to chemical energy via acyclic or cyclic electron flow but the uphill transport of energy from PSI to PSII.

One mechanism by which the uphill transport can be accounted for is anti-Stokes fluorescence, that ascertains the existence of a luminescence center that absorbs a low-energy photon, and then emits a high-energy photon the excess is supplied by the thermal absorption from the optical medium and this results in cooling. The Anti Stokes cooling was first proposed by Pringsheim (Pringsheim, 1929) and was contradicted by (Landau, 1946). However there have been some recent works (Djeu and Whitney, 1981, Qin et al., 2006) confirming Anti Stokes cooling due to anti-Stokes fluorescent cooling in a single center (ASFCSC) as well as by energy transfer (ASFCET) among centers. If Anti Stokes cooling is observed in plants the scenario is likely to mimic the ASFCET. An entirely different mode of uphill energy transport that is tightly coupled to the downhill transport of energy may be the two photon process. It was suggested by Fong (Fong, 1975) that photoionization of active chlorophylls may involve the summation of two red excitations. Similar suggestion of two photon process in biological process was given by (Hallenbeck and Benemann, 2002). It may be noted that there is a long history of development of two photon process and recently the use in microscopic process is well known (Diaspro et al., 2005).

### Thermal implications of coupling between PSI and PSII

Thermodynamics is a topic that prevails across the boundary of classical and quantum description of nature (Prigogine, 1978). Irrespective of whether the uphill energy transport is mediated by anti Stokes fuorescence or by two red photon process or by another biological process that results in a similar outcome, we can derive certain important implications. Positive dynamic efficiency of the coupled process would imply absorption of thermal energy from the surroundings.

Let us assume a phenomenological approach (Onsager and Machlup, 1953). The driving force in such system *X* can be identified as entropy change (Dasgupta, 2014) Δ*S_V_* for emission at frequency *ν*. The flux terms *J* in the photon capture process emerges as 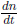, *n* being the number of photons emitted during an emission process. If we reserve the superscripts 1 and 2 to represent the PSI and PSII system, the phenomenological equations that follow are:

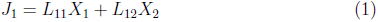

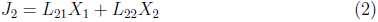

In (1)–(2) *L* is the Onsager matrix showing the symmetry *L*_12_ = *L*_21_. We thus express emission rates from (1) and (2) by equations (1–2). The rate of intensity change from PSI (J1) as a linear combination of excitation due to PSI and PSII and so is for PSII. If we further assume that the entropy associated with non-radiative changes are given by:

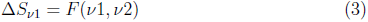

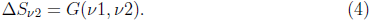

where we assume that the nonradiative exchange at emission for the PSI and PSII are functions *F* (*ν*1, *ν*2) and *G*(*ν*1, *ν*2), the coupling between PSI, PSII (independent of whether they are classical or quantum) being ultimately expressible by the system:

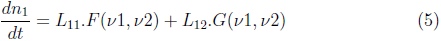

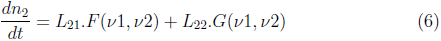

Eventually, the equation (6) represents a coupled dynamical system which may show oscillation depending on the functional dependence of *F* and *G* on *ν*1 and *ν*2 as reported by (Sen et al., 2014). The functional nature in fact depends on the transitional probability between the different states and the rate of mutual energy transfer between the same.

The efficiency of the non-equilibrium coupled process is given by:

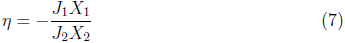

where, the LHS of the equation (3) represents the efficiency of the energy transfer between PSI and PSII. In equation (7) ‘2’ represents the downhill the energy transfer and therefore the driving force and ’1’ is the uphill transfer that is driven.

Let us now examine whether the efficiency of photosynthetic machinery defined by equation (7) can be near unity. It follows that if *η* = 1 − *ϵ* where ϵ → 0, leads to a situation det |*L*| = 0, a condition that is not permitted by second law of thermodynamics. . Eventually, the photon energy of a median energy photon at 600nm is 2.07 eV, and for 8 moles of such photons the energy absorbed is 381 Kcal. It takes 114 Kcal to reduce one mole of CO2 to hexose, so the theoretical efficiency is 114/381 or 30% (Lea, Drake, 1999). It thus follows that the efficiency of energy transfer looked upon theoretically or from simple experimental consideration is much less than one. The question is what the thermodynamic fate of the unavailable energy remains. In other words what follows is:

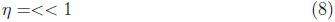

We can rewrite equation (7) as:

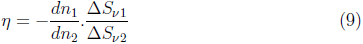

If we assume that the energy transfer is an one photon process, 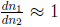 the inequality that is satisfied by the coupled process is given by, 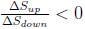, the term in subscript representing the total change in entropy in the uphill and downhill process. What follows is that the coupled process can operate only if there are opposite signs of entropy change in the coupling steps. If we further assume the entropy enthalpy compensation (Battistuzzi et al., 2004, Chodera and Mobley, 2013) the change in enthalpy in the respective processes will also have opposite sign. Expressed in simpler words, a thermodynamic coupling between the photosynthetic centres would demand one of the steps to to be endothermic and the other exothermic, the latter recognizable as the downhill process from PSII to PSI. The PSI to PSII transition would be evidently endothermic in nature, and whenever the integrated transition involves PSI to PSII, that is there is an enhancement of the PSII to PSI ratio, on expects a predominance of the cooling process.

## Materials and methods

### Isolation of Thylakoid Membranes

Dark-adapted leaves rinsed in distilled water were homogenized with ice-cold homogenization buffer in a pre-chilled mortar pestle (5 ml of homogenization buffer were added per gm of leaf). The slurry was then filtered through two layers of cheese cloth. The pellet was gently suspended in washing buffer (1 ml washing buffer per gm of tissue) and the suspension was left in ice for 10 minutes. The suspension was then centrifuged at 2800g for 5 minutes at 4°*C*. The washing was repeated twice and the pellet was suspended in storage buffer and total chlorophyll was estimated. Aliquots of thylakoid membranes were snap cooled in liquid N2 and stored at −80°*C*. The preparations were protected from light and kept ice-cold during the isolation procedure. Chlorophyll was extracted in 80% aqueous acetone and determined according to Hipkins and Baker (1986).

### Plant Material

Arachis hypogea plants were grown in a growth chamber at 25C with a 16 hour photoperiod under 40*μEm*^−2^*S*^−1^ of irradiance. Leaflets of an average diameter of 1 cm were collected from 3-4 week old plants.

### Treatments of Arachis Thylakoids

Thylakoids were subjected to DCMU (10*μM*), Nigericin (6*μM*), (GNP) Gold nanoparticle (Singha et al., 2010), (AgNP)Silver nanoparticle (Roy et al., 2010) and chlorophyll nanoparticle (containing mixture with approximate composition 163nM chlorophyll A 126 nM chlorophyll B) treatments for 20 minutes before subjecting them to irradiance of 200*μmol* of photons *m*^−2^*S*^−1^ intensity at 25C.

### Gold nanoparticle (GNP) synthesis

The synthesis was carried out using an accepted method (Frens, 1973) modified using the method reported by (Singha et al., 2010). An aqueous solution of HAuCl4 (*20mM, 25mL*) was brought nearly to boiling condition and stirred continuously with a stirrer. Freshly prepared tri-sodium citrate solution (38.8mM) was added quickly at a time. The citrate concentration is related to the particle size, resulting in a change in color from pale yellow to deep red. The temperature was brought down to normal and colloidal solution was stirred for an additional few minutes and the volume makeup was performed with excess citrate. Typical plasmonic responance for AuNP was at 530nm, which according to standard literature corresponds to a size close to 40nm, final atomic concentration of gold being 200*μM*.

### Silver nanoparticle (AgNP) synthesis

The silver nanoparticles were synthesized by reducing silver nitrate 20 mM (AgNO3) solution in presence of sodium borohydrate (100mM). Citrate acts as a stabilizer in the preparation. The stirring was performed at normal temperature leading to a change in color from transparent to yellow rendering AgNP formation. After 15 minutes of stirring at room temperature the prepared AgNP were incubated in ice for half an hour before use. The plasmon resonance was observed in the range 420nm, that corresponds to a size 40nm. The final atomic concentration of silver is 200*μM*.

### Organic chlorophyll nanoparticle (ChlNP) synthesis

Chlorophyll is the main pigment biomolecule of the photosynthetic machinery. In polar solvents like water hydrophobic tetrapyrrole moiety of the chlorophyll molecule tends to aggregate among them securing their hydrophobic core by π – π stacking interaction. Size controlled nanoparticle formulation from this porphyrin molecule is evident in appropriate capping and stabilizing conditions. The stabilizer prevents agglomeration and is a factor for hydrodynamic property determination. Water soluble ChlNP were synthesized and stabilized by prolonged sonication of spinach chlorophyll composite using citrate as a stabilizing and capping agent.

### Dynamic light scattering (DLS)

The hydrodynamic properties of the nano structures were characterized by DLS instrument. Study were undertaken using a Malvern Nano ZS80 (UK) dynamic light scattering set up equipped with a 532nm excitation laser source.

### 77K Fluorescence Measurement (Krause and Weis, 1984)

Thylakoid (400*μgChl/ml*) samples were subjected to irradiances of various intensities for various time periods over duration of 5 hrs as described above. Following irradiance treatment samples were diluted in a medium containing 70% glycerol (w/v), 20 mM HEPES buffer, pH 7.8, 5 mM MgCl2, and 0.33 M sorbitol to a final concentration of 10*μgChl/ml* before freezing them immediately in liquid nitrogen. Low temperature fluorescence emission spectra were recorded using a Hitachi 4050 Spectroflurimeter (Japan) where the excitation light was provided from a Xenon light source. The excitation monochromator was set to 480 nm (band width of 5 nm). Emission was scanned from 650 to 800 nm (band width of 3 nm). 77K fluorescence spectra were normalized at 722 nm fluorescence and ratios for *F_PSI_/F_PSII_* were calculated and plotted against corresponding time of exposure to irradiance. The nano structure influence on the energy transfer dynamics had also been investigated. Three sets of data were acquired from each of two independent isolations.

### Thermal Imaging Technique

Thylakoid samples were treated with 200*μEm*−^2^*S*^−1^light and thermal image was monitored for indicated time period. Representative temperature profile is shown in four petri plates. Snapshots were acquired with an interval of 10 minutes with the of FLIR IR camera. The thermal data analysis is carried out following the procedure described earlier (Azharuddin et al., 2014) using FLIR software and MATLAB 14.0 ( Mathworks USA).

### Temperature Measurement with Thermocouple

Temperature of differently treated thylakoid samples was monitored with 5*SRTC* – *TT* – *T* – 30 – 36 thermocouple connected with *HH*147*U* Four-Channel Handheld Data Logger from Omega Engineering, during with 200*μEm*^−2^*S*^−1^ light treatment for indicated time period.

## Results

Table 1 summaries the essential points put forward in this paper. The third column of the table connects the spectral changes and the heating pattern obtained. As the storage buffer is always held as control, the heating and cooling in the post illumination state is always evaluated relative to the storage buffer (see the first row of Table 1). This is confirmed both by the thermal imaging measurement and the thermo couple measurement.

**Table 1:** Coupled energy transfer and thermal output of the photosynthetic machinery in presence of nano scale and physical perturbation by protonophores and LEF blocker DCMU.

| Sample under illumination | Perturbant | Thermal and Spectral |
| --- | --- | --- |
| Storage Buffer | (-) | (-) |
| Thylakoid membrane | (-) | Cooling PSI PSII |
| Thylakoid membrane | GNP (23nm) | Heating PSI(0) PSII(0) |
| Thylakoid membrane | ChlNP (86nm) | Heating PSI(++) PSII(+) |
| Thylakoid membrane | SNP (97nm) | Cooling PSI(+) PSII(++) |
| Thylakoid membrane | Protonophore (Nigericin, CCCP) | Heating PSI(+) PSII(0) |
| Thylakoid membrane | DCMU | PSI(0) PSII(0) |

### Monitoring the Temperature of Differently Treated Thylakoid Samples during Irradiance Treatment

Thylakoid solutions of 400*μg/ml* concentration were suspended in storage buffer and irradiated with 200*μEm*^−2^*S*^−1^ light. Equal volume of storage buffer incapable of photosynthetic light reaction served as negative control. Thermal imaging was done during the course of white light treatment for 3 hours (Fig. 1A) to check the temperature of the thylakoid solution. The thermal image of each sample was collected for the time period of 3 hours with an interval of 10 minutes by acquiring video for 2mins with the of FLIR IR camera. The maximum, minimum and average temperature of selected zone was extracted out from the videos by help of FLIR software. We found continuous cooling of thylakoid sample with respect to buffer (Fig 1B). At the end of 3 hours we noticed thylakoid solution is cooler by 2^O^*C* than buffer solution. We performed the same experiment for another 1.5 hrs and measured the temperature of the solution with thermocouple which is direct contact method for measuring temperature (Fig. 1C). A similar 2^O^*C* cooling is observed at the end of 1.5 hrs (Fig 1D).

**Figure 1:**
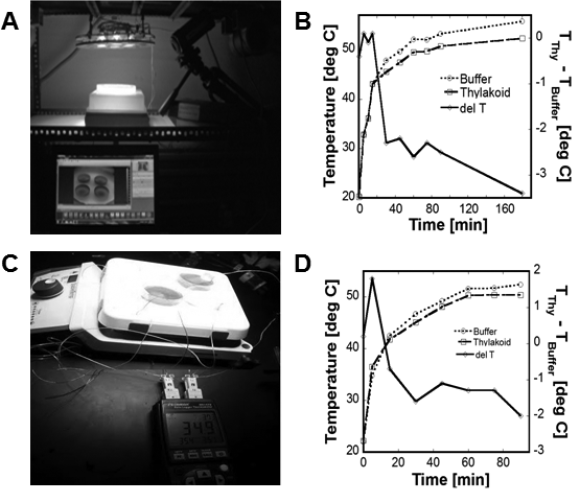
Cooling of the thylakoid membrane with respect to the storage buffer. The experiments were performed using thermal imaging (upper panels A,B) and thermo-couple(lower panels C,D). Thylakoid and storage buffer are taken in two petri plate and treated with 200*μEm*^−2^*S*^−1^ light for 4 hr and the temperature profiles of the respective measurements thermal imaging(panel B) and thermocouple(panel D) are shown

### Effect of inhibitors on Cooling Process

Diuron or DCMU (3-(3,4-dichlorophenyl)-1,1-dimethylurea) interrupts the photosynthetic electron transport chain in photosynthesis. In this experiment our objective was to study the effect of this inhibitor on cooling process to confirm that observed cooling is related to photosynthetic electron transport. We have taken thylakoid solution of 400*μgml*^−1^Chl. Concentration and DCMU treated thylakoid of same Chl concentration. Storage buffer serves as control. These samples are irradiated with 200*μEm*^−2^*S*^−1^ light as earlier and temperature was measured with thermal imaging camera. We have noticed the cooling of thylakoid solution with respect to the controls as earlier but the DCMU treated thylakoid shows no cooling and behaves as control storage buffer at the end of 3 hrs of light treatment (Fig 2). We checked the role of Δ*pH* on generation and maintenance of cooling in thylakoid by using uncouplers like nigericin and CCCP. These uncouplers operate by electro-neutral dissipation of Δ*pH* by accelerating non-phosphorylating *H*+ fluxes across thylakoids. Thylakoid sample was incubated with 50mg/ml Nigericin and 5 mg/ml CCCP for 20 minutes and then the temperature was monitored for 1 hour. After 1 hours thylakoid sample treated with Nigericin and CCCP was 2°*C* hotter than control thylakoid (data not shown).

**Figure 2:**
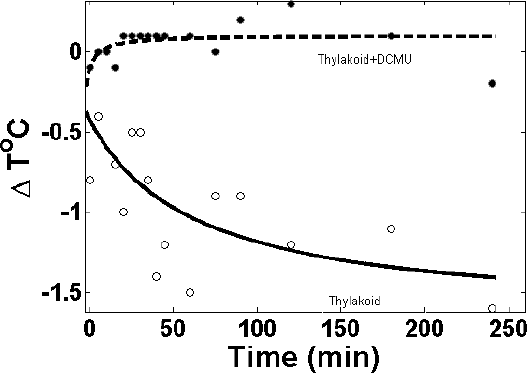
Effect of photosynthetic inhibitor on cooling process. Photosynthetic linear electron flow blocker DCMU (dotted line) on the thermal output of the thylakoid membrane,solid line representing the control thylakoid membrane (similar effects seen with other uncouplers (data not shown)

### Effect of Perturbation Agents on Cooling Process

We incubated thylakoid sample with perturbation agents like (a) Gold nanoparticle, GNP, (b) Silver nanoparticle, AgNP (c) Chlorophyll nanoparticle, ChlNP for 20 minutes and then temperature of thylakoid sample with FLIR. Thermal imaging was done during the course of white light treatment for 2 hours. Different temperature profile was noted for different nanoparticle treated thylakoid sample (Fig. 3). At the end of 4 hour thylakoid sample treated with GNP showed a little higher temperature than control thylakoid (Fig. 4A). This difference in temperature was more prominent in case of ChlNP treated sample, which was showing almost 1.7^o^C difference (Fig. 4C). But thylakoid sample treated with AgNP showed 1.9^o^ C lower temperature than control thylakoid (Fig. 4E). We monitored the energy transfer between PSII and PSI in otherwise similar condition as temperature measurement by *F_PSI_*/*F_PSII_* calculation at 77K. With GNP treatment *F_PSI_*/*F_PSII_* was > 1 indicating PSII to PSI energy transfer (Fig. 4B). Similarly with ChlNP treatment *F_PSI_*/*F_PSII_* was >> 1, indicating PSII to PSI energy transfer to a greater extent (Fig. 4D). On the other hand treatment with AgNP resulted in *F_PSI_*/*F_PSII_* was << 1 indicating PSI to PSII energy transfer which is energetically unfavorable (Fig. 4F).

**Figure 3:**
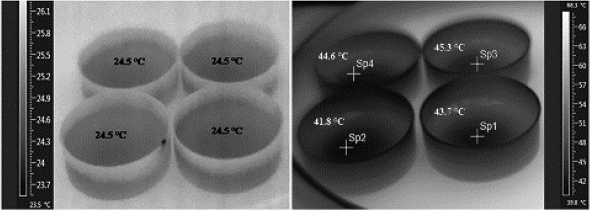
Effect of perturbation agent on cooling process. Left panel represents the nanoparticle administered petri-plates with photosynthetic membranes;right panel shows representative images of the same after prolong white light illumination. In this panel (right) Sp1 represents control thylakoid membrane, Sp2 represents SNP treated membrane, Sp3 represents ChlNP treated and Sp4 represents GNP treated thylakoid membrane fractions. The side bar represents the color code for temperature

**Figure 4:**
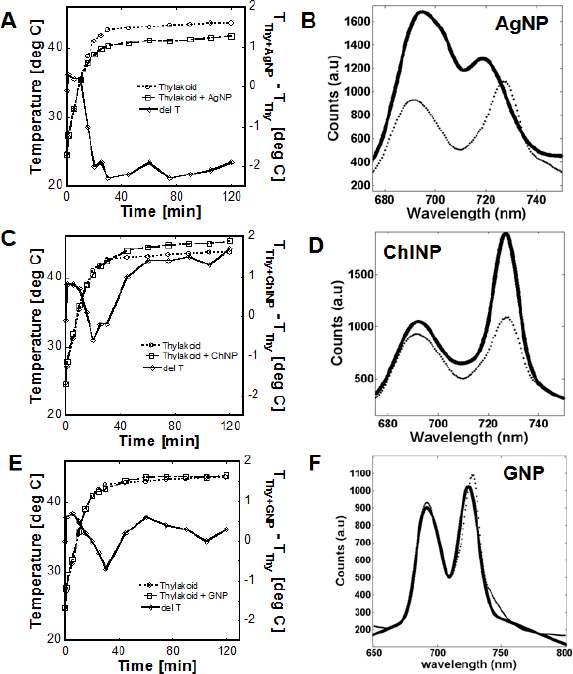
Correspondence between Thermal and Spectral Profiles: Correlated changes in thermal profiles (panels A,C,E) and corresponding spectral profiles (panels B,D,F) in presence of AgNP, ChlNP and AuNP are shown. Lighter and darker lines representing the presence and absence of nanoparticles. The spectral profiles were obtained from 77K fluorescence of the thylakoid membrane suspension with or without the nano particle treatement as described in the methods section. Cooling corresponds to increase in PSI/PSII ratio (also see Table 1)

## Discussions

The urban heat island (UHI) phenomenon is now a well known effect, though the subject often get masked by socio-political debates. Cooling effect by plant species are normally explained by presence of two factors namely, blocking of sunlight (Keith et al., 2010) and secondly, by evapo-transpiration(Penuelas et al., 2009, Pokorny, 2001). The present analysis suggests that blocking of Sunlight actually can elevate the UHI phenomena in a counterintutive manner as the anti-stokes cooling will be spared. The emergence of urban heat island (Akbari, 2009) is known for some time and the introduction of green roofs has been emphasized. Recent studies indicate that New york city and New Delhi can be 10^o^F hotter than the outlying area. The difference can be partially accounted for by the combined effect of evapo-transpiration, and by our analysis by uphill electron transport and the resultant anti Stokes cooling. The other arguments often given is that reflection of Sunlight is the main mechanism by which the cooling occurs, dark color leaves causing more cooling than light colored ones. But the matter that is often disregarded is that the amount reflected does not fully account for the cooling by trees, the amount of light that fluoresces as a result of absorption is also important. In most of the plants systems the coupling between PSI and PSII leads to further complications, the uphill transport leading to cooling as we have shown in the introduction. While the Stokes emission is likely to cause heating like reflection, the uphill energy transport does exist and the residual cooling that may follow may highlight a hitherto ignored aspect of the cooling phenomena. The study opens up a new question why certain plants are more effective in cooling than other plants.

With these general observations in mind, two points deserves some special mention. This is the inhibition of cooling by electron flow blockers like DCMU (Fig. 2), and by other uncouplers (data not shown). The other point is the effect of nanoparticles that interferes with the heating/cooling process. The nanoparticles used in this work have special spectral properties in the range that has an overlap with the excitation spectra of the photocentres (peak at 480nm). While the AgNP behaves as a cut-off filter, absorbing and 420, facilitating specific 480 excitation of the PSI or II, the GNP, with 530nm actually cuts off the 480nm absorbtion partially. Although, both are composed of noble metals,this difference in optical characters may lead to the reverse thermodynamic effects (heating and cooling). The reversed changes in the 77K spectra (Fig. 4) also support this inference. We do not understand the exact mechanism of insignificant heating induced by gold nanoparticles,and cooling by the silver nanoparticle, but the plasmonic argument we have put forward, provides a rational base for the probable mechanism underlying the reported effects.

At this stage of this work, the ChlNP effect can be considered as a phenomenological observation. It may be noted that had cooling been based on light absorption alone, the result obtained by Chl nanoparticle should have been the reverse. The global relation that we find satisfied for each nanoparticle is that the increase in PSII/PSI ratio causes cooling(AgNP) the reverse causes heating(ChlNP),and no significant change in the ratio (GNP) causes no significant heating or cooling.

Thus we find cooling as result of coupled uphill and downhill energy transport and its artificially manipulation by nanoparticles. Agents causing higher uphill energy flow from PSI to PSII (i.e enhancement of PSII/PSI ratio, see Table 1) always causes cooling and the reverse is also true. Thus the minimal information that we can extract from the above data is that cooling is indeed due to coupling between the two photo centres and the cooling can be artificially manipulated by spectrally active nanomaterials. Some implications of the result may be of practical importance, e.g., how how this cooling impacts the plant ecology and how the phenomenon can be exploited for energy saving, or how the essential thermodynamic principle can be used in temperature controlled phyto-filtration (Nash, 2003)and in other similar green technology ventures.

